# Multiplexed single-cell imaging reveals diverging subpopulations with distinct senescence phenotypes during long-term senescence induction

**DOI:** 10.1101/2024.10.14.618296

**Authors:** Garrett A. Sessions, Madeline V. Loops, Brian O. Diekman, Jeremy E. Purvis

**Affiliations:** Department of Cell Biology and Physiology, The University of North Carolina at Chapel Hill, Chapel Hill, NC 27599, Unites States of America; Department of Biology, The University of North Carolina at Chapel Hill, Chapel Hill, NC 27599, Unites States of America; Department of Molecular Genetics and Cell Biology, The University of Chicago, Chicago, IL 60637, United States of America; Thurston Arthritis Research Center, The University of North Carolina at Chapel Hill, Chapel Hill, NC 27599, United States of America; Joint Department of Biomedical Engineering, University of North Carolina at Chapel Hill, Chapel Hill, North Carolina 27599 and North Carolina State University, Raleigh, North Carolina 27695, United States of America; Lineberger Comprehensive Cancer Center, University of North Carolina at Chapel Hill, North Carolina 27599, United States of America; Department of Genetics, The University of North Carolina at Chapel Hill, Chapel Hill, NC 27599, United States of America; Computational Medicine Program, The University of North Carolina at Chapel Hill, Chapel Hill, NC 27599, United States of America

**Author notes:** **Corresponding authors:** Jeremy Purvis, Department of Genetics, The University of North Carolina School of Medicine, 11018C Mary Ellen Jones, Campus Box #7264, Chapel Hill, NC 27599-7264.

**Keywords:** Cellular senescence, Senescence-associated secretory phenotype (SASP), Senescence Induction, DNA Damage

## Abstract

Cellular senescence is a phenotypic state that contributes to the progression of age-related disease through secretion of pro-inflammatory factors known as the senescence associated secretory phenotype (SASP). Understanding the process by which healthy cells become senescent and develop SASP factors is critical for improving the identification of senescent cells and, ultimately, understanding tissue dysfunction. Here, we reveal how the duration of cellular stress modulates the SASP in distinct subpopulations of senescent cells. We used multiplex, single-cell imaging to build a proteomic map of senescence induction in human epithelial cells induced to senescence over the course of 31 days. We map how the expression of SASP proteins increases alongside other known senescence markers such as p53, p21, and p16^INK4a^. The aggregated population of cells responded to etoposide with an accumulation of stress response factors over the first 11 days, followed by a plateau in most proteins. At the single-cell level, however, we identified two distinct senescence cell populations, one defined primarily by larger nuclear area and the second by higher protein concentrations. Trajectory inference suggested that cells took one of two discrete molecular paths from unperturbed healthy cells, through a common transitional subpopulation, and ending at the discrete terminal senescence phenotypes. Our results underscore the importance of using single-cell proteomics to identify the mechanistic pathways governing the transition from senescence induction to a mature state of senescence characterized by the SASP.

## Introduction

Aging drives an ever-increasing load on healthcare systems, with more than half of the global disease burden resulting from age-related diseases[1]. This burden can be expected to rise as the average age of the global population continues to increase. Cellular senescence is a phenotypic state associated with diseases that emerge at high rates with aging, such as osteoarthritis[2–4], neurodegenerative disorders[5,6], and macular degeneration[7]. Through the production and secretion of pro-inflammatory factors known as the Senescence Associated Secretory Phenotype (SASP), even a small number of senescent cells can have an outsized effect on the tissue environment[8]. Thus, identifying and eliminating senescent cells is an appealing target for therapeutics aimed at reducing the burden of age-related disease[9,10]. Senescence can be induced both *in vivo* and *in vitro* by subjecting healthy cells to long-term stresses such as oxidative stress[3] and DNA damage[11,12]. However, it is unclear how long cells must be exposed to each stress to induce senescence and SASP factors, or what sequence of molecular states they undergo en route from health to senescence[13].

There is tremendous value in understanding how cells respond to stress, succumb to it, and what happens after cells succumb to that stress. A major challenge is that senescence is a highly heterogeneous phenomenon[14] and one clear source of heterogeneity is temporal. There is strong evidence that key features of senescence are dynamic[13,15,16], but many studies focus on a single timepoint and thus the conclusions reflect only a snapshot of the senescence phenotype. Thus, it is currently unknown to what degree the observed heterogeneity in the field can be explained by differences in timepoint selection between experiments. By combining a granular time course experiment with the multidimensional proteomics analysis, we hope to expand on the role of temporal heterogeneity as a contributing factor to the overall heterogeneity of senescence.

Subpopulation heterogeneity is another source of complexity when trying to interpret data from aggregate analysis. The process of senescence induction is unlikely to be uniform across the individual cells within a culture, but single-cell analysis provides the capacity to identify subpopulations of senescent cells with distinct molecular features. Further, approaches to define the trajectories of cells over pseudotime can give insight into cells that are at different points along the path towards these various senescence subpopulations within a given experiment. Other systems, such as studies of apoptosis, have successfully mapped both the temporal and subpopulation heterogeneity of those processes[17]. A better understanding of the contributions of temporal and subpopulation heterogeneity in senescence will provide a more accurate and useful framework for understanding the role of senescence in age-related disease.

Here, we reveal the temporal dynamics of senescence induction in a cell type with well-characterized cell cycle dynamics, retinal pigment epithelial (RPE) cells[18,19]. We use a novel proteomics technique, iterative indirect immunofluorescence imaging[20] (4i), to profile induced senescent cells across a month of continuous senescence induction. We find that much of the observed heterogeneity of cellular senescence can be explained by the gradual, long-term changes in senescence phenotype as well as the emergence of discrete populations of senescent cells with different molecular signatures over those longer timeframes. Age-related disease is the result of a lifetime of accumulated damage; thus, we anticipate that the senescence phenotype “matures” in response to this damage over long timeframes. By understanding how the senescence phenotype alters with increasing depths and durations of senescence, we will be better equipped to identify potential targets for intervention.

## Materials and Methods

### Cell lines and culture conditions and treatments

Retinal pigment epithelial cells (hTERT RPE-1, ATCC, CRL-4000) were used for all experiments. RPE cells were cultured at 37 °C and 5% CO_2_ in DMEM (Glibco, 11995-065) with 10% fetal bovine serum (FBS; Sigma, TMS-013-B), GlutaMAX (Glibco, 35050-061), and penicillin/streptomycin (P/S; ThermoFisher Scientific 15140148). To induce senescence, 1 µM of etoposide (MedChem, HY-13629/CS-1774) was included at each feed (MWF) and left on the cells between feeds.

### Antibodies

Antibodies used in this study (**Supplemental Table 1**) were either previously selected using BenchSci and tested in prior work[18,19] or selected for relevance to senescence and tested prior to inclusion. Testing of antibodies was performed to ensure correct staining above background and that antibodies could be eluted using the established 4i protocols. Hoechst (Sigma-Aldrich 33258) was used as a nuclear stain. The following secondary antibodies were used as appropriate – Alexa Fluor 647 Donkey anti-goat (Invitrogen A21447), Alexa Fluor 568 Donkey anti-mouse (Invitrogen A10037), Alexa Fluor 568 Donkey anti-rabbit (Invitrogen A10042), Alexa Fluor 488 Donkey anti-rabbit (Invitrogen A32790).

### Iterative Indirect Immunofluorescence Imaging (4i)

Samples were prepared as previously described[18–20]. Stitched 5×5 images were collected using a Nikon Ti2 Eclipse inverted microscope using a Plan Apo LambdaD 20x objective lens (NA=0.8) with a Teledyne Photometrics Kinetix sCMOS camera. Image acquisition performed using the following filer cubes: DAPI (Semrock DAPI-3060A), AF488 (Semrock GFP-4050B), AF568 (Semrock mCherry-C), AF647 (Semrock LED- Cy5-5070A). Image acquisition and stitching was performed using NIS-Elements HCA JOBS software to enable the automated imaging of the entire well and plate. Images from each round of 4i were aligned in Python 3.7 using the StackReg library and manually checked for accurate alignment. Segmentation was performed in Python 3.7 using the CellPose library and feature quantification and extraction was performed using the region properties library of Scikit-image.

### Data visualization and trajectory inference

Data were visualized in Python 3.7 using Potential of Heat-diffusion for Affinity-based Transition Embedding[21] (PHATE) in three dimensions. PHATE was performed on z-normalized data that had been subsampled to 1200 cells per timepoint using Sketch[22]. Trajectory inference was performed in R using the Slingshot package with PHATE coordinates and anchored on the ground truth population of cells not treated with etoposide. Trajectories from Slingshot were overlaid on PHATE plots in Python 3.7. All other visualizations were performed using Python 3.7 and Jupyter Notebooks using the environments and notebooks hosted at (https://github.com/PurvisLabTeam/publication_code_repo) under the papers title in the folder labeled analysis_code.

### Data availability and code availability

Single cell datasets are available here (https://github.com/PurvisLabTeam/publication_code_repo) under the papers title in the folder labeled source_data. Code used for processing 4i datasets is available here (https://github.com/PurvisLabTeam/4i_pipeline).

## Results

### Long-term senescence induction

We used a well-established model of the human cell cycle[18,19], retinal pigment epithelial (RPE) cells, to investigate the temporal dynamics of senescence induction. RPE cells were continuously exposed to 1 µM etoposide, a Topoisomerase II inhibitor that induces DNA damage, for up to 31 days **(Fig 1A)**. We profiled cells by 4i for 16 markers of senescence, SASP, and cell cycle signaling **(Supp Table 1)** at 12 time intervals over 31 days after initiating induction with etoposide plus an untreated control condition. The collected images were processed, screened for artifacts, and quantified to produce a tabular dataset **(Fig 1B)**. As cells were subjected to increasing durations of etoposide treatment, they began to take on visual markers of senescence such as increased size and disrupted morphology **(Supp Fig 1)**.

**Figure 1.**
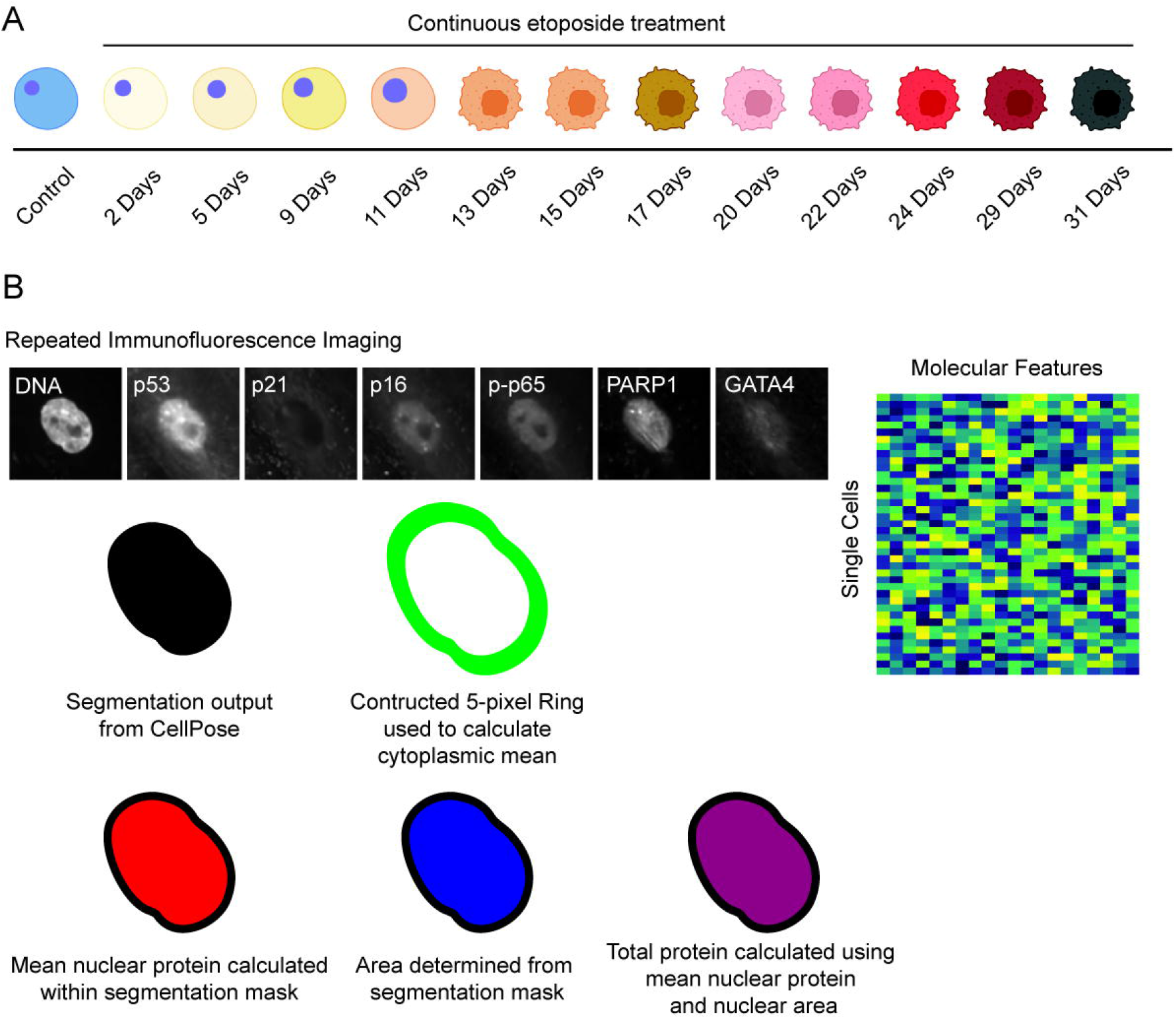
Experimental design and data processing pipeline. A. Experimental timepoints collected for 4i image analysis. B. Schematic for quantifying features from single-cell images. Nuclear area, shown in blue determines the area used to calculate the nuclear mean protein levels and the total nuclear protein levels. A 5-pixel ring is constructed around the nucleus to determine the mean cytoplasmic protein levels. Multiple image features for each cell are used to produce a tabular data matrix for downstream analysis. Cells in panel A were created with Biorender.

### Aggregate population analysis

To gain an overview of senescence induction over the 31-day time course, we first visualized aggregate changes in total nuclear (total), mean nuclear (mean), and mean cytoplasmic (cytoplasmic) protein for the entire population of cells **(Fig 1B)**. This analysis revealed two major phases of changes in senescence associated factors: an induction phase ranging from the day 0 control until day 11; and a steady state phase ranging from day 11 until the final timepoint at day 31. This trend was observed most strikingly when calculating the overall increase in nuclear area **(Fig 2A)**. The consistent increase in size until day 11 and maintenance of that size until day 31 can be observed in representative images as **(Fig 2B)**. Across all 16 proteins analyzed in this study, total protein levels closely mirrored the temporal pattern established for nuclear area **(Fig 2A)**, suggesting that the increase in nuclear area is a critical event that facilitates increased total protein abundance **(Supp Fig 2)**. Total protein dynamics for 6 key proteins are shown in **Fig 2C**. Established senescence markers such as p53, p21, and p16 followed previously reported dynamical trends[2,23–25] at the bulk total protein level. For example, persistent DNA damage provoked through continuous application of etoposide drove an early upregulation of total p53 protein. Similarly, total levels of p21 and p16 increased at each time point from day 2 through day 11 before reaching a plateau phase. In addition to these DNA damage and cell cycle regulators, we observed temporal changes in proteins involved in inflammatory signaling. For example, total phosphorylated p65 (p-p65), a key component of the NF-KB pathway, was upregulated in lockstep with total p53. Finally, at the end of the induction phase at day 11, the transition to a steady state phase was accompanied by switch-like increases in the total abundances of GATA4 and PARP1 (**Supp Fig 3)**.

**Figure 2.**
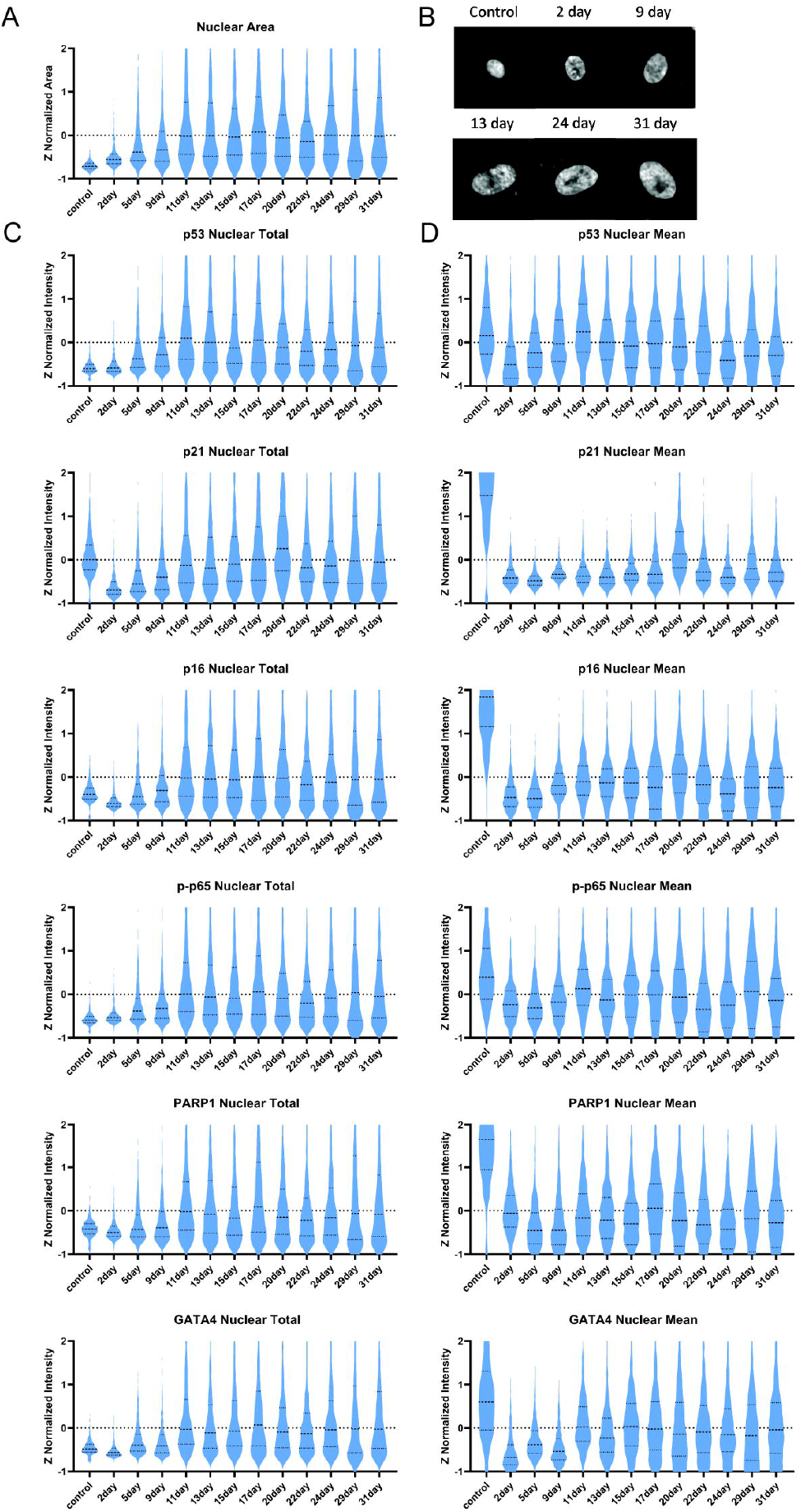
Visualization of key features across all collected timepoints. A) Nuclear area increases at the bulk level until day 11. B) Representative images of the increasing nuclear size across timepoints. C) Total protein is determined by multiplying the area in pixels and the nuclear mean intensity of a given feature for each cell. Viewed in bulk, total protein increases as nuclear area increases, leveling off at day 11. D) Nuclear mean quantifies the protein concentration of each cell. Featured proteins all observed a decline in nuclear mean as nuclear area increases in bulk.

Unlike the total protein levels, mean protein concentration decreased rapidly in critical senescence proteins, with some recovery towards control levels over time. Mean levels of p53, p16, p-p65, PARP1, and GATA4 declined from control to day 2 then began to increase in concentration until the 11-day timepoint. None of these proteins except p53 returned to the mean levels observed in the control cells. Mean levels of p21 declined rapidly and did not meaningfully increase at any subsequent timepoint (**Fig 2D)**. These dynamics are due in part to the rapid increase in nuclear area upon etoposide treatment, which dilutes all mean protein concentrations.

In summary, time series of aggregate single-cell distributions shows an initial response phase over the first 11 days characterized by accumulation multiple stress and inflammatory signaling proteins, followed by a stable period of response with apparently fewer changes in protein levels. We next sought to fully utilize the single-cell measurements to detect the presence of potential subpopulations of healthy and senescent cells in this aggregated cell population.

### Identifying subpopulations of healthy and senescence cells

We next explored whether cells clustered into different subpopulations and what transitions they underwent while progressing from an unperturbed, healthy state to these discrete subpopulations. To visualize all cells in their various molecular states, which are defined by their unique combinations of protein levels, we used the nonlinear dimensionality reduction method PHATE[21] (Potential of Heat-diffusion for Affinity-based Transition Embedding) to embed the high dimensional data into three dimensions. In parallel, we performed *k*-means clustering on the entire dataset to identify distinct subpopulations. To choose the number of clusters in a principled way, we chose the number of clusters that maximized the percentage of unperturbed cells in a single cluster, which we referred to as the healthy cells. This strategy produced 10 clusters that could be sorted into six cell-type subpopulations that included the healthy cells, three “terminal” clusters, and two transitional populations **(Fig. 3A)**. Here, terminal clusters represent groups of cells with a large molecular distance from the healthy cells. Two of the terminal clusters expressed high levels of known senescence markers but differed in terms of nuclear area and the concentration of senescence associated and SASP proteins. The third terminal cluster responded to etoposide that made them distinct from the healthy cells, but this cluster did not express high total levels or high concentrations of senescence markers. In addition, one of the transitional populations mapped onto the PHATE structure in between the healthy cells and the two senescence clusters, whereas the second transitional population occupied the space between the healthy cells and the non-senescent terminal cluster. Because we are specifically focused on the transition to senescence, in the subsequent analysis we primarily focused on the healthy cells, the two senescent terminal clusters, and the single senescence transitional cluster that presumably gave rise to the two senescence subpopulations.

**Figure 3.**
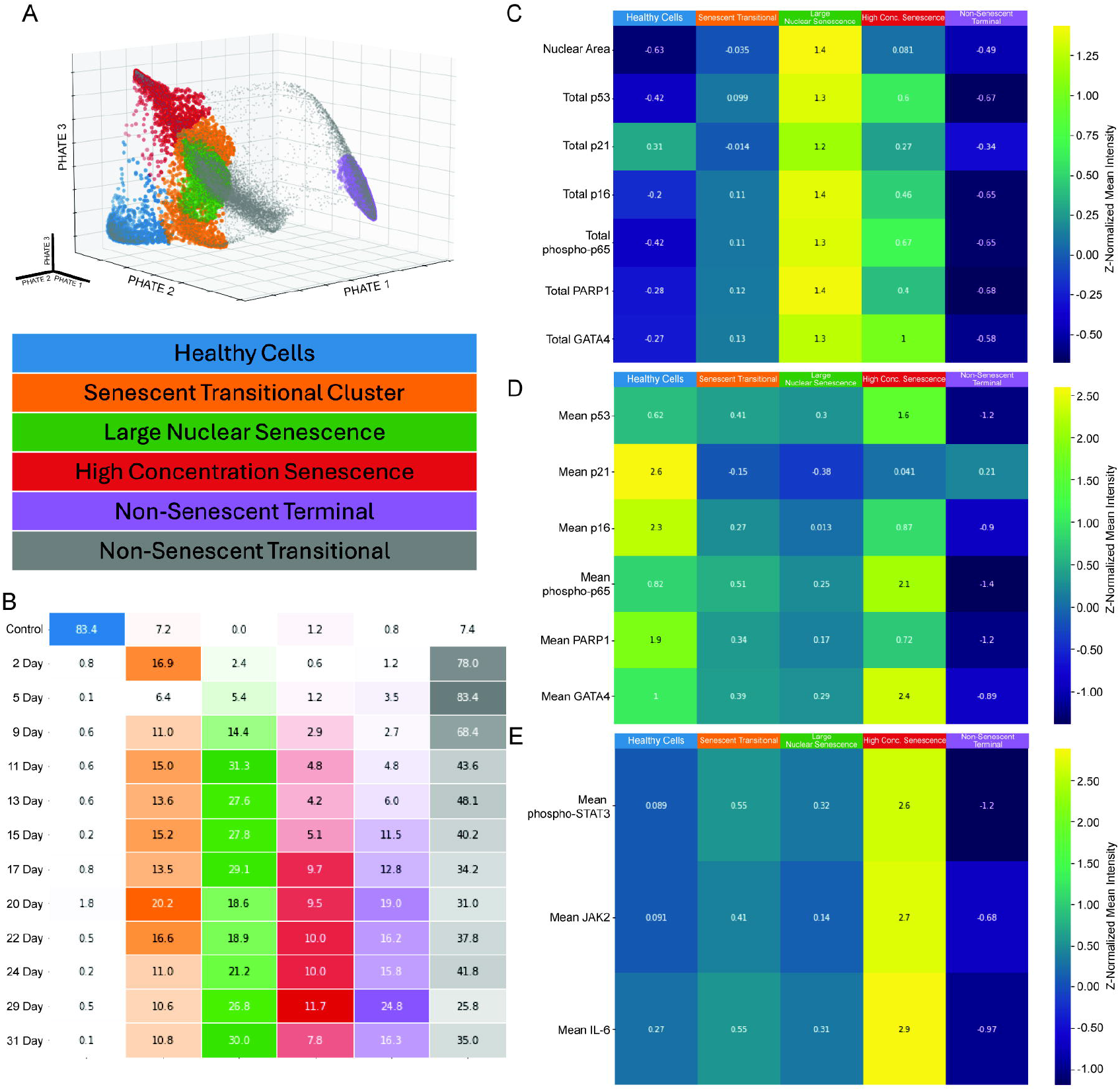
Key clusters are differentiated by nuclear area and mean protein in the nucleus and cytoplasm. A) The five key subpopulations projected onto the PHATE structure. B) Distribution of cells at each timepoint across the five clusters of interest and the non-senescent transitional clusters. C) Nuclear area and total protein for each cluster of interest. D) Nuclear mean protein for each cluster of interest. E) Cytoplasmic mean protein for select IL-6 pathway proteins.

To understand how these subpopulations arise in response to etoposide, we first quantified the contribution of each subpopulation at individual time points. This analysis shows that the transitional subpopulation arises immediately, with the senescence transitional cluster accounting for nearly 17% of the cells at day 2 (**Fig 3B**). By contrast, the three terminal clusters did not accumulate to a significant extent until much later in the time course, with the large nuclear senescence cluster arising strongly at day 11 and the high concentration senescent cell population not arising strongly until day 17. Thus, what appeared in the aggregate data to be a single stable phase is revealed to be a composition of multiple subpopulations which accumulate cells at different rates. The transitional cluster identified through *k*-means clustering is superficially similar to the high concentration senescent cell cluster by nuclear area and total nuclear protein levels of key senescence proteins (**Fig 3C**). However, the transitional cluster lacks the increased protein concentrations of the high concentration senescence cluster, indicating that it is less active for core senescence functions (**Fig 3D**). Additionally, the transitional cluster lacks cytoplasmic concentrations of IL-6 pathway proteins observed in the high concentration senescence cluster (**Fig 3E**).

### Inferring the temporal pathway from healthy to senescent cells

The PHATE plots and time point representation for each cluster provide some indication of the temporal order in which the identified populations emerge during the response to etoposide. To gain additional understanding of the dynamics of senescence induction, we used Slingshot[26] to infer the temporal trajectory of molecular states that cells pass through en route from healthy to senescence (**Fig 4A)**. In this analysis we identified three lineage paths, each originating in the healthy cell cluster and ending in one of three distinct terminal clusters: non-senescent terminal, large nuclear senescence, and high-concentration senescence. These trajectories produce a pseudotime axis which describes how a hypothetical cell may pass through the molecular states captured by the trajectory. We next asked how individual molecular features changed as cells moved along each individual trajectory. We used locally estimated scatterplot smoothing (Loess) curves to calculate the mean and variation of each feature along the trendline. For example, **Fig 4B** shows how nuclear area—a key senescence feature—varies among individual cells in the entire population, and **Fig 4C** shows how nuclear area changes along each lineage trajectory. Only one lineage, the large nuclear senescence lineage, showed substantial changes in nuclear size along its trajectory. In contrast, the high-concentration senescence lineage showed a slight increase in nuclear size that eventually leveled off. The non-senescent terminal lineage results in distinctly altered cells that also showed a transient increase in nuclear area, but which are distinct from the two senescence lineages in that it lacks expression of common senescence markers such as p53, p21, and p16.

**Figure 4.**
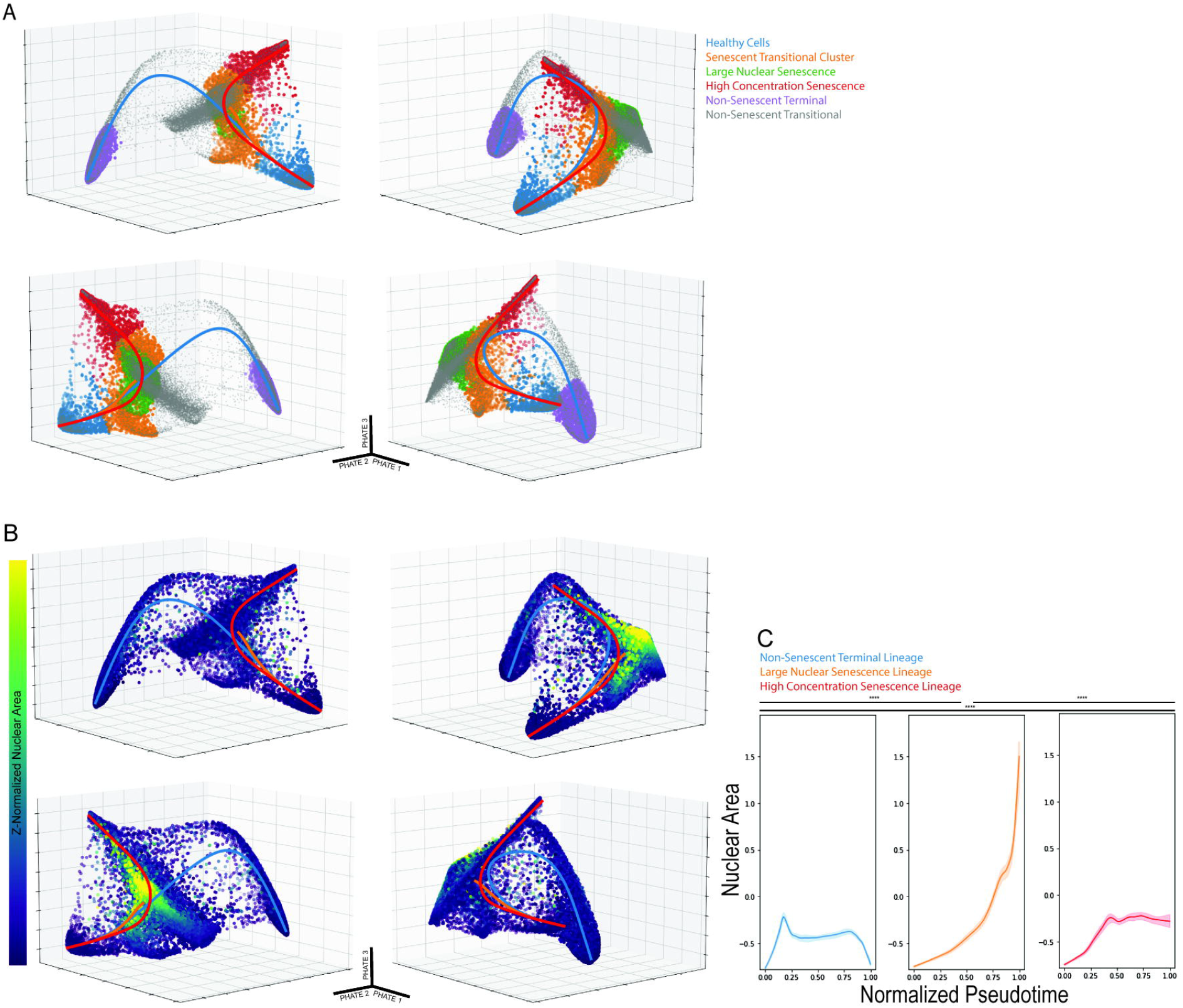
3D PHATE and K means clustering identify discrete populations of interest. Lineage tracing through these clustere populations shows three distinct paths from health to senescence. A) Five biologically meaningful clusters, highlighted here, projected onto the 3D PHATE structure. Slingshot produces three trajectories rooted in the healthy cell cluster, which all pas through the senescent transitional cluster, ending in three distinct terminal clusters. B) Z-normalized nuclear area projected the 3D PHATE structure alongside Slingshot Trajectories. C) Nuclear area plotted against the pseudotime axis of each of th trajectories. The Kolmogorov-Smirnov test was applied to test the differences between the plotted curves. (* = p<0.05, ** = p<1e-100, *** = p<1e-200, **** = p<1e-300)

### Protein dynamics in lower dimensional embedding

To gain a more detailed view of how healthy cells transition toward each of the three terminal clusters—large nuclear senescence, high-concentration senescence, and non-senescent terminal—we examined total protein levels for 6 key senescence features within each subpopulation (**Fig 5A**) and then traced the temporal dynamics of these features along each lineage trajectory (**Fig 5B**). We found that p53, p-p65, PARP1, and GATA4 levels increased uniformly across pseudotime in the large nuclear senescence lineage. p16 underwent a transient decline like that seen in the aggregate analysis before increasing. p21 declined in total protein abundance until the cell exited the transitional cluster and entered the terminal large nuclear senescence cluster, at which point the abundance of p21 rapidly increased. Total levels of all six of the key proteins was strongly correlated with nuclear area, with the large nuclear senescent cell lineage having statistically significantly higher total protein levels in the final 10% of the pseudotime when compared to the high concentration senescent cell lineage (**Fig 5C**). Comparison of changes in mean protein levels also suggested that the switch-like behavior observed in the aggregate timepoint analysis of GATA4 and PARP1 (**Fig 2C**) is largely driven by the increased total nuclear protein of the large nuclear senescent cells. In contrast to total protein levels, the mean nuclear protein levels of key senescence proteins accumulated in different regions of the PHATE structure and showed a greater variety of trends **(Fig 5D)**. Regardless of the lineage, the early pseudotime is characterized by a decline in mean protein concentration. However, once the high concentration senescence population exited the transitional cluster and entered the high concentration terminal cluster, we observe that mean protein levels rose beyond initial levels in the cases of p53, p-p65, and GATA4 **(Fig 5E,F)**. Notably, the high concentration senescent cells showed sharp increases in mean levels of GATA4 and PARP1 while these factors decreased in concentration in the large nuclear senescent cells **(Fig 5E,F**). Taken together, these results suggest rapid accumulation of protein concentration in distinct subpopulations of senescent cells that are masked by aggregate cell analysis.

**Figure 5.**
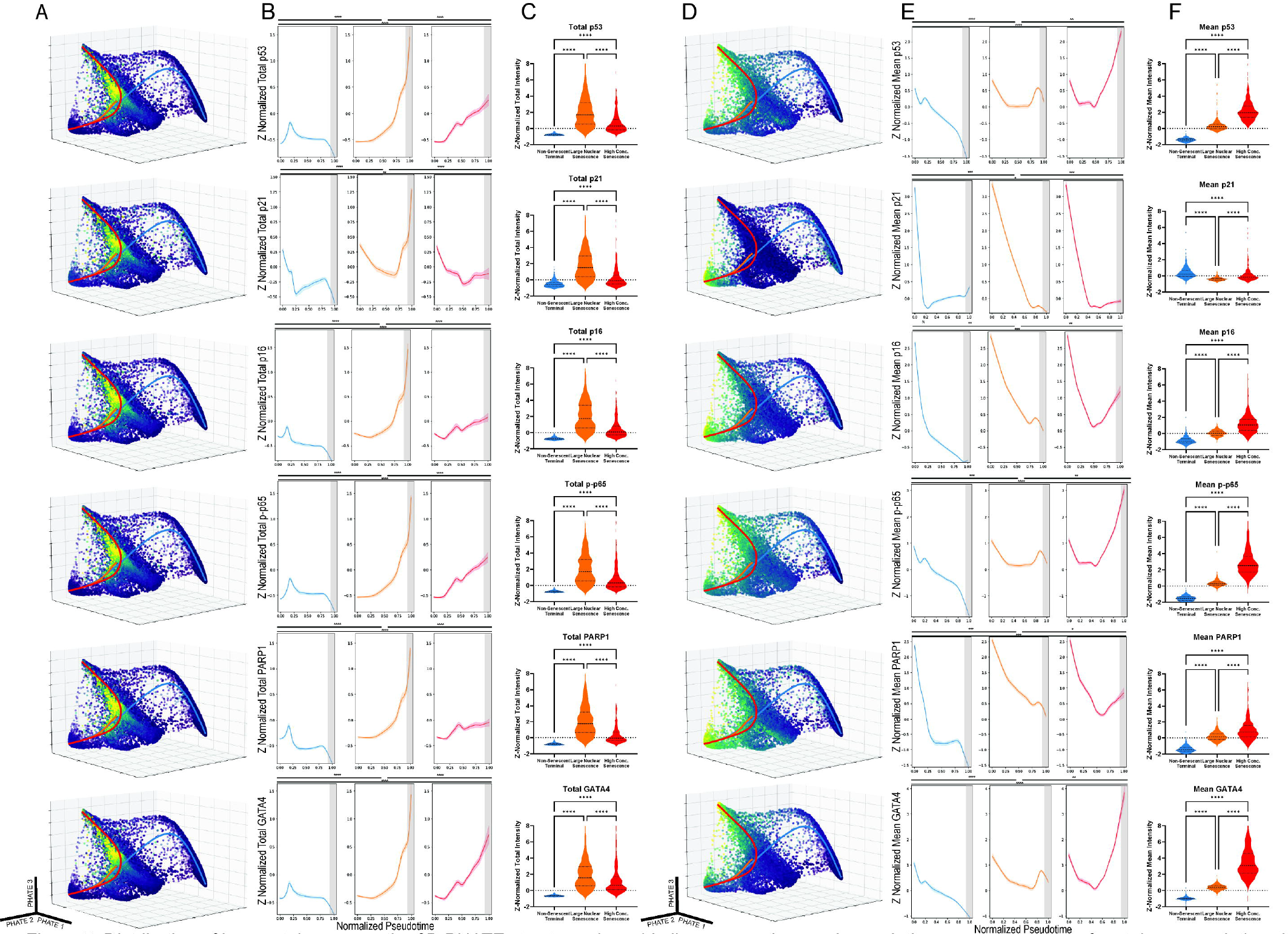
Distribution of key proteins across the 30 PHATE structure alongside lineage overlays and pseudotime representations of protein accumulation. A) Distribution of total nuclear protein across the 30 PHATE structure highlighting six key proteins. B) Total nuclear protein accumulation plotted against the pseudotime of each lineage. The last 10% of each lineage is highlighted in gray. The Kolmogorov-Smirnov test was applied to test the differences between the plotted curves.(*= p<0.05, **= p<1e-100, ***= p<1e-200, ****= p<1e-300) C) Comparison of the total protein levels in the originating healthy cell cluster, the large nuclear senescence cluster, and the high concentration senescence cluster. D) Mean nuclear protein accumulation plotted against the pseudotime of each lineage. The last 10% of each lineage is highlighted in gray. The Kolmogorov-Smirnov test was applied to test the differences between the plotted curves.(*= p<0.05, **= p<1e-100, ***= p<1e-200, ****= p<1e-300) E) Mean nuclear protein accumulation plotted against the pseudotime of each lineage. F) Comparison of the Mean protein levels in the originating healthy cell cluster, the large nuclear senescence cluster, and the high concentration senescence cluster.

### Differential regulation of the SASP in distinct subpopulations of senescent cells

We next examined the potential functional consequences of the molecular differences observed between the large nuclear and high concentration senescent cell populations. Examination of IL-6 pathway proteins in the cytoplasm showed that the high concentration senescent cells are the primary producers of IL-6, a common SASP factor. JAK2 protein levels also accumulate strongly along the high concentration lineage **(Fig 6A)**. This trend was further supported by the Loess curves plotted using the Slingshot-derived pseudotime axis for JAK2 **(Fig 6B)**. Notably, this trend was not obvious from analysis of the aggregate data of all cells at each timepoint (**Fig 6C)**. pSTAT3, another key component of the IL-6 pathway, was similarly regulated, and again the trends shown by trajectory analysis were not observable in the aggregate data **(Fig 6D-F)**. The differences in IL-6 expression itself were similarly striking, with the large nuclear senescent cells showing no meaningful changes from controls and the high concentration senescent cells showing a drastic increase in cytoplasmic IL-6 (**Fig 6G-I**). These three components of the IL-6 pathway co-occur in individual cells and are upregulated in lockstep with each other. This lockstep regulation is evidence for autocrine reinforcement of the high concentration phenotype.

**Figure 6.**
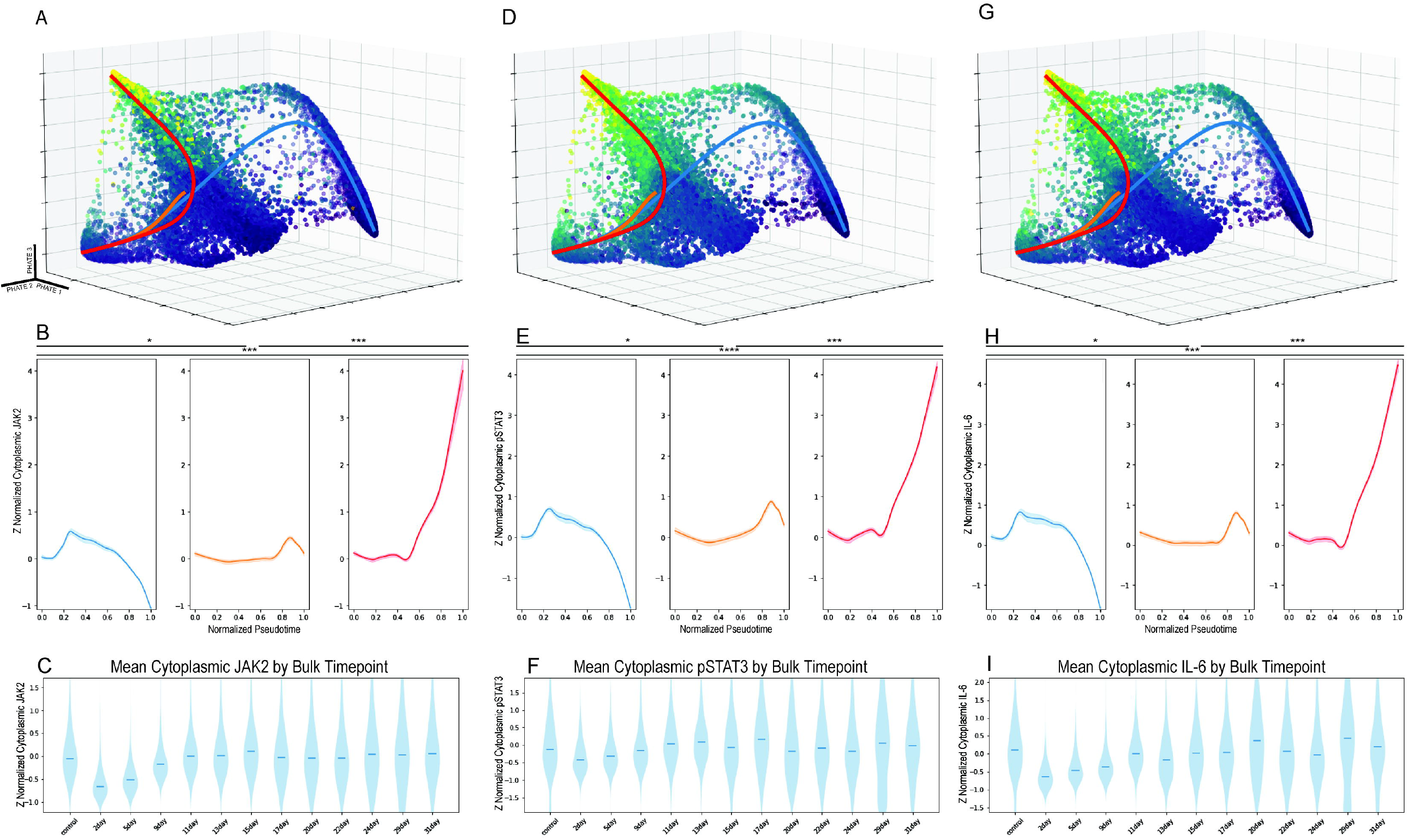
Mean cytoplasmic protein accumulation for IL-6 pathway proteins. A) Distribution of mean cytoplasmic JAK2 across the 3D PHATE structure B) Mean cytoplasmic JAK2 accumulation plotted against the pseudotime of each lineage. (* = p<0.05, ** = p<1e-100, *** = p<1e-200, **** = p<1e-300) C) Cytoplasmic mean of JAK2 pooled by timepoints. D) Distribution of mean cytoplasmic pSTAT3 across the 3D PHATE structure E) Mean cytoplasmic pSTAT3 accumulation plotted against the pseudotime of each lineage. (* = p<0.05, ** = p<1e-100, *** = p<1e-200, **** = p<1e-300) F) Cytoplasmic mean of pSTAT3 pooled by timepoints. G) Distribution of mean cytoplasmic IL-6across the 3D PHATE structure H) Mean cytoplasmic IL-6accumulation plotted against the pseudotime of each lineage. (* = p<0.05, ** = p<1e-100, *** = p<1e-200, **** = p<1e-300) I) Cytoplasmic mean of IL-6 pooled by timepoints.

## Discussion

How senescence-associated proteins are regulated over time, and among individual cells, is a complex process. This work highlights the critical need for comprehensive analysis of relevant proteins and pathways across time and at the single-cell level when attempting to understand how senescence arises in healthy cell populations. We found two discrete pathways of senescence induction: one present in cells with large nuclei and high total protein levels of canonical senescence markers that support cell cycle arrest, and one present in cells with a high concentration of senescence proteins that appears to support SASP production with an emphasis on the IL-6 pathway (**Fig 7**). Crosstalk between these two systems could be reinforced by switch-like behavior in transcriptional regulators like GATA4 and PARP1, which reinforce cell cycle arrest and SASP production through orthogonal pathways.

**Figure 7.**
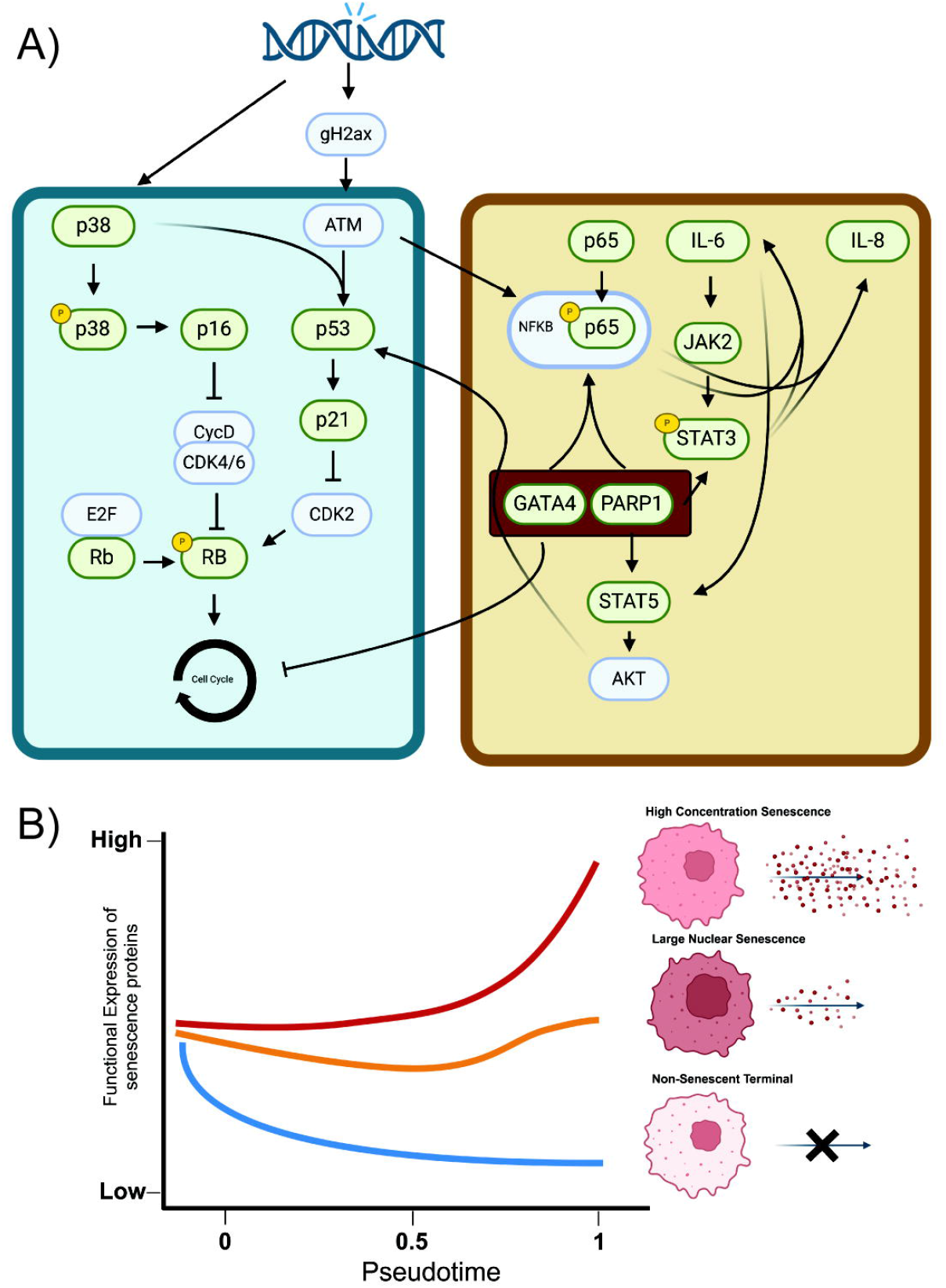
Schematic of proposed protein interactions. A) Green boxes are proteins directly measured in this dataset. Blue boxes are proteins inferred from observed interactions. GATA4 and PARP1 highlighted in red to emphasize their potential role in regulating the switch to metabolically active small nuclear senescent cells. B) Three overall patterns emerge from the single-cell analysis. High concentration senescent cells increase mean expression of senescence associated proteins and SASP factors as pseudotime increases. Large nuclear senescent cells show a very limited increase or no increase at all. Non-senescent terminal cells show an extreme decline in senescence associated proteins and SASP factors. Both panels were created with Biorender.

Cells under etoposide stress rapidly exit the set of molecular states associated with healthy cells and begin to accumulate in either the transitional cluster or, after a time delay, one of the three terminal clusters. Once etoposide stress has been applied, the transitional cluster is always occupied by some fraction of the total cell population. This indicates that cells may be continually entering or exiting this transitional molecular state.

Given that all three trajectories pass through this cluster, we infer that it represents a transitional state in which cells have not yet entered a molecular state that will push them towards one of the three terminal clusters.

Molecular markers indicate that these cells have not mounted a DNA damage response and have no detectable SASP component (**Fig 3**).

From this senescence transitional cluster, three discrete terminal populations emerge, two of which have hallmarks of senescence. Nuclear morphology and size as well as overall cell size[27–29] are often one of the initial indicators of senescence. Indeed, one of the three lineages/populations derived from the multiplexed dataset is defined in part by very large nuclei. These large nuclear senescent cells have increased total nuclear protein levels for most measured features. However, the mean protein levels of these cells tend to stay stable or reduce in concentration. It is possible that the increased protein levels seen in these cells, which would have a large effect on non-single cell, aggregate analyses, is driven by the increasing cell size as a compensatory mechanism to maintain a stable protein concentration.

In contrast, there is a second lineage/population of senescent cells that arises which is not defined by increased nuclear area, but by increased concentration of senescence-associated proteins. These cells still have larger nuclei than unperturbed control cells, but their primary defining features appear to be increased concentrations of proteins such as p53, phospho-p65, and other markers of senescence and the SASP.

Trajectory analysis shows that this population arises later than the large nuclear senescence population (**Fig 5**) and may be responsible for the bulk of the SASP features measured in this work. Beyond the difference in nuclear protein concentration, the large nuclear and high concentration senescent cell populations exhibit key differences in their ability to support SASP secretion, inferred here by the cytoplasmic concentration of IL-6 pathway proteins. High concentration senescent cells appear to be upregulating protein synthesis in response to the DNA damage stress, increasing the concentration of key senescence proteins to active downstream responses like the IL-6 pathway. DNA damage has been previously shown to induce pro-inflammatory SASP[30,31], making this pathway an ideal target for study. The cytoplasmic concentration of JAK2, phospho- STAT3 and IL-6 are all greatly increased in the high concentration senescent cells. We infer from this that the rate of secretion of IL-6 out of the cell is higher as well. Future work will determine to what extent the SASP factors in this subpopulation may affect other cells through paracrine mechanisms.

The third population of cells appears to represent a heterogeneous mix of non-senescent cells that express higher levels of damage than the unperturbed populations. This population emerges from the senescent transitional cluster and passes into the non-senescent transitional cluster before ending in the non-senescent terminal cluster. This population is ill-defined by the panel of markers used in this work, as those markers were chosen to identify senescent cells and to understand the SASP. As a result, we cannot speak at length to specific features of this population, other than to say that these cells occupy a distinct molecular state, different from both healthy cells and from both populations of senescent cells. It is possible that these cells are dysfunctional in meaningful ways that are the result of paracrine influences by etoposide-induced senescent cells. It is also possible that they are cells that have resolved the stress of etoposide treatment in a way that does not result in senescence.

In conclusion, our single-cell analysis revealed that the temporal dynamics of senescence induction is a key contributor to the observed heterogeneity of cellular senescence. That is, cells reach different states at different rates such that any instantaneous measurement of aggregate cells will reveal a different mix of distinct subpopulations. The two key populations explored here—the large nuclear senescent and the high concentration senescent—do not arise in great number until day 11 or day 17 respectively (**Fig 3**), although a small number of cells exhibiting these characteristic molecular states can be detected at even the very first timepoint. In this study, the analysis of IL-6 pathway components highlights the weaknesses of only measuring senescence features as an aggregate of many cells. The majority of the IL-6 pathway proteins were expressed at meaningful levels only in the high concentration senescence population (**Fig 6B,E,H)**, which comprised just 7.8% of all cells in the day 31 timepoint **(Fig 3B)**. The aggregate analysis missed the contribution of this population entirely, showing only the steady stabilization of IL-6 pathway proteins after day 11 **(Fig 6C,F,I)**.

Thus, had we only measured the aggregate data, averaging out many cells into a single measurement and combining a range of distinct molecular states, we would have masked a key temporal event and the discrete populations that arise in response to a continuous direct DNA damage challenge. By combining a highly multiplexed set of single cell protein measures and a detailed, granular time course experiment, our work suggests that senescent cells undergo a series of time-dependent interactions between previously described senescence markers. Importantly, our work reveals large differences in SASP potential between discrete molecular states of senescent cells that could be easily missed when combined with sizable populations of non-senescent cells that persist and even increases after a full month of etoposide treatment.

From a therapeutic perspective, the removal or modulation of senescent cells by drug treatment is an appealing target for addressing the rising incidence of age-related disease. The senescence transitional state represents an appealing target for future work, as there may be molecular switches or other mechanisms present at this stage that could allow for the prediction of which terminal state the cell will ultimately enter.

Resolving these mechanisms in detail could provide novel molecular targets for therapeutics. For example, GATA4 may be a potential target for reducing the entry of transitional cells into a SASP-producing program exhibited by the high concentration senescent cells. The emergence of GATA4 and PARP1 as transcriptional regulators that potentially stabilize the senescence phenotype highlights promising targets for potential therapeutics. In particular, the concentration of GATA4 is markedly increased in the high concentration senescent cells. There are several roles for GATA4 in senescence indicated in the literature, but the most interesting in regard to these findings is its role in the reinforcement of cell cycle arrest and the NFκB pathway[32]. Future work targeting the inhibition of GATA4 could produce a potential senomorphic target, valuable for its reduction of the SASP. The earlier these changes can be detected in individual cells, the less time the tissue environment will be exposed to harmful SASP.

## Acknowledgements

We thank Dr. Sam Wolff, Dr. Phil Coryell, Dr. Richard Loeser, Dr. Kasia Kedziora, and Lorenzo Santarina. This work was supported by grants R01-AG081734 (B.O.D.), NSF- 2242980 (J.E.P.), R01-GM138834 (J.E.P.), and R01-CA280482 (J.E.P.).

## Statements and Declarations

The authors declare that they have no financial or non-financial conflicts of interests.

## Notes

https://github.com/PurvisLabTeam/publication_code_repo

https://github.com/PurvisLabTeam/4i_pipeline

### Competing Interest Statement

The authors have declared no competing interest.

## References

[1] Chang AY, Skirbekk VF, Tyrovolas S, Kassebaum NJ, Dieleman JL. Measuring population ageing: an analysis of the Global Burden of Disease Study 2017. Lancet Public Health 2019;4:e159–67. 10.1016/S2468-2667(19)30019-2.

[2] Diekman BO, Sessions GA, Collins JA, Knecht AK, Strum SL, Mitin NK, et al. Expression of p16 ^INK 4a^ is a biomarker of chondrocyte aging but does not cause osteoarthritis. Aging Cell 2018;17:e12771. 10.1111/acel.12771.

[3] Coryell PR, Diekman BO, Loeser RF. Mechanisms and therapeutic implications of cellular senescence in osteoarthritis. Nat Rev Rheumatol 2021;17:47–57. 10.1038/s41584-020-00533-7.

[4] Jeon OH, David N, Campisi J, Elisseeff JH. Senescent cells and osteoarthritis: a painful connection. J Clin Invest 2018;128:1229–37. 10.1172/jci95147.

[5] Cohen J, Torres C. Astrocyte senescence: Evidence and significance. Aging Cell 2019;18:e12937. 10.1111/acel.12937.

[6] Bhat R, Crowe EP, Bitto A, Moh M, Katsetos CD, Garcia FU, et al. Astrocyte Senescence as a Component of Alzheimer’s Disease. PLoS ONE 2012;7:e45069. 10.1371/journal.pone.0045069.

[7] Lacour M, Kiilgaard JF, Nissen MH. Age-Related Macular Degeneration: Epidemiology and Optimal Treatment. Drugs Aging 2002;19:101–33. 10.2165/00002512-200219020-00003.

[8] Basisty N, Kale A, Jeon OH, Kuehnemann C, Payne T, Rao C, et al. A proteomic atlas of senescence-associated secretomes for aging biomarker development. PLOS Biol 2020;18:e3000599. 10.1371/journal.pbio.3000599.

[9] Childs BG, Gluscevic M, Baker DJ, Laberge R-M, Marquess D, Dananberg J, et al. Senescent cells: an emerging target for diseases of ageing. Nat Rev Drug Discov 2017;16:718–35. 10.1038/nrd.2017.116.

[10] Chaib S, Tchkonia T, Kirkland JL. Cellular senescence and senolytics: the path to the clinic. Nat Med 2022;28:1556–68. 10.1038/s41591-022-01923-y.

[11] te Poele RH, Okorokov AL, Jardine L, Cummings J, Joel SP. DNA damage is able to induce senescence in tumor cells in vitro and in vivo. Cancer Res 2002;62:1876–83.

[12] Ogrodnik M, Salmonowicz H, Gladyshev VN. Integrating cellular senescence with the concept of damage accumulation in aging: Relevance for clearance of senescent cells. Aging Cell 2019;18:e12841. 10.1111/acel.12841.

[13] De Cecco M, Ito T, Petrashen AP, Elias AE, Skvir NJ, Criscione SW, et al. L1 drives IFN in senescent cells and promotes age-associated inflammation. Nature 2019;566:73–8. 10.1038/s41586-018-0784-9.

[14] Bitencourt TC, Vargas JE, Silva AO, Fraga LR, Filippi-Chiela E. Subcellular structure, heterogeneity, and plasticity of senescent cells. Aging Cell 2024:e14154. 10.1111/acel.14154.

[15] Hernandez-Segura A, De Jong TV, Melov S, Guryev V, Campisi J, Demaria M. Unmasking Transcriptional Heterogeneity in Senescent Cells. Curr Biol 2017;27:2652-2660.e4. 10.1016/j.cub.2017.07.033.

[16] Wechter N, Rossi M, Anerillas C, Tsitsipatis D, Piao Y, Fan J, et al. Single-cell transcriptomic analysis uncovers diverse and dynamic senescent cell populations. Aging 2023. 10.18632/aging.204666.

[17] Spencer SL, Sorger PK. Measuring and Modeling Apoptosis in Single Cells. Cell 2011;144:926–39. 10.1016/j.cell.2011.03.002.

[18] Stallaert W, Kedziora KM, Taylor CD, Zikry TM, Ranek JS, Sobon HK, et al. The structure of the human cell cycle. Cell Syst 2022;13:230-240.e3. 10.1016/j.cels.2021.10.007.

[19] Stallaert W, Taylor SR, Kedziora KM, Taylor CD, Sobon HK, Young CL, et al. The molecular architecture of cell cycle arrest. Mol Syst Biol 2022;18. 10.15252/msb.202211087.

[20] Gut G, Herrmann MD, Pelkmans L. Multiplexed protein maps link subcellular organization to cellular states. Science 2018;361:eaar7042. 10.1126/science.aar7042.

[21] Moon KR, Van Dijk D, Wang Z, Gigante S, Burkhardt DB, Chen WS, et al. Visualizing structure and transitions in high-dimensional biological data. Nat Biotechnol 2019;37:1482–92. 10.1038/s41587-019-0336-3.

[22] Baskaran VA, Ranek J, Shan S, Stanley N, Oliva JB. Distribution-based sketching of single-cell samples. Proc. 13th ACM Int. Conf. Bioinforma. Comput. Biol. Health Inform., Northbrook Illinois: ACM; 2022, p. 1– 10. 10.1145/3535508.3545539.

[23] Serrano M, Lin AW, McCurrach ME, Beach D, Lowe SW. Oncogenic ras Provokes Premature Cell Senescence Associated with Accumulation of p53 and p16INK4a. Cell 1997;88:593–602. 10.1016/S0092-8674(00)81902-9.

[24] Krishnamurthy J, Torrice C, Ramsey MR, Kovalev GI, Al-Regaiey K, Su L, et al. Ink4a/Arf expression is a biomarker of aging. J Clin Invest 2004;114:1299–307. 10.1172/JCI22475.

[25] Ressler S, Bartkova J, Niederegger H, Bartek J, Scharffetter-Kochanek K, Jansen Dürr P, et al. p16 ^INK4A^ is a robust in vivo biomarker of cellular aging in human skin. Aging Cell 2006;5:379–89. 10.1111/j.1474-9726.2006.00231.x.

[26] Street K, Risso D, Fletcher RB, Das D, Ngai J, Yosef N, et al. Slingshot: cell lineage and pseudotime inference for single-cell transcriptomics. BMC Genomics 2018;19:477. 10.1186/s12864-018-4772-0.

[27] Heckenbach I, Mkrtchyan GV, Ezra MB, Bakula D, Madsen JS, Nielsen MH, et al. Nuclear morphology is a deep learning biomarker of cellular senescence. Nat Aging 2022;2:742–55. 10.1038/s43587-022-00263-3.

[28] Sherwood SW, Rush D, Ellsworth JL, Schimke RT. Defining cellular senescence in IMR-90 cells: a flow cytometric analysis. Proc Natl Acad Sci 1988;85:9086–90. 10.1073/pnas.85.23.9086.

[29] Wiley CD, Campisi J. From Ancient Pathways to Aging Cells—Connecting Metabolism and Cellular Senescence. Cell Metab 2016;23:1013–21. 10.1016/j.cmet.2016.05.010.

[30] Rodier F, Coppé J-P, Patil CK, Hoeijmakers WAM, Muñoz DP, Raza SR, et al. Persistent DNA damage signalling triggers senescence-associated inflammatory cytokine secretion. Nat Cell Biol 2009;11:973–9. 10.1038/ncb1909.

[31] Tchkonia T, Zhu Y, Van Deursen J, Campisi J, Kirkland JL. Cellular senescence and the senescent secretory phenotype: therapeutic opportunities. J Clin Invest 2013;123:966–72. 10.1172/JCI64098.

[32] Kang C, Xu Q, Martin TD, Li MZ, Demaria M, Aron L, et al. The DNA damage response induces inflammation and senescence by inhibiting autophagy of GATA4. Science 2015;349:aaa5612. 10.1126/science.aaa5612.

