## Supplementary figures and images for "Multiplexed single-cell imaging reveals diverging subpopulations with distinct senescence phenotypes during long-term senescence induction"

### Supplemental Figure 1

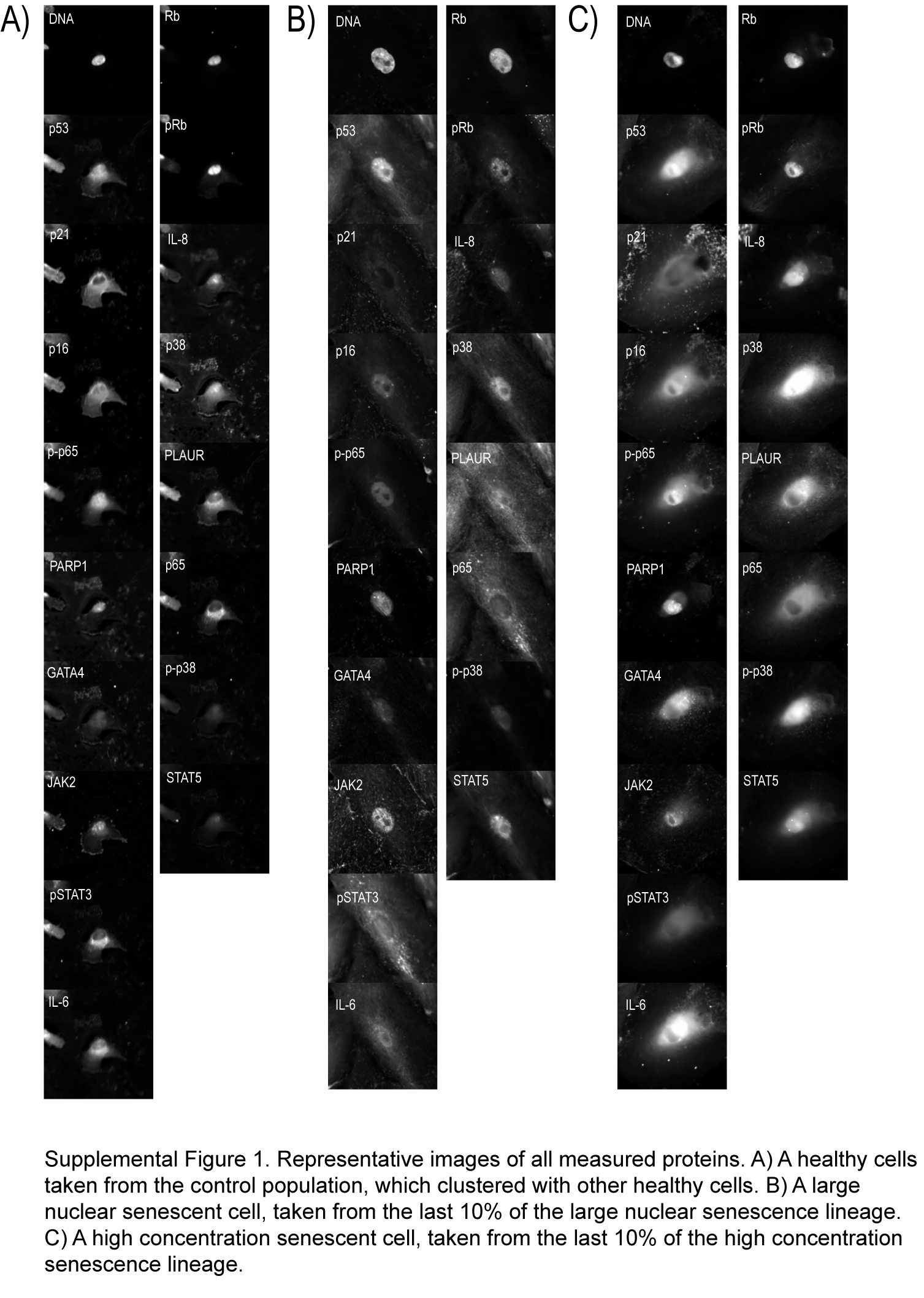

### Supplemental Figure 2.1

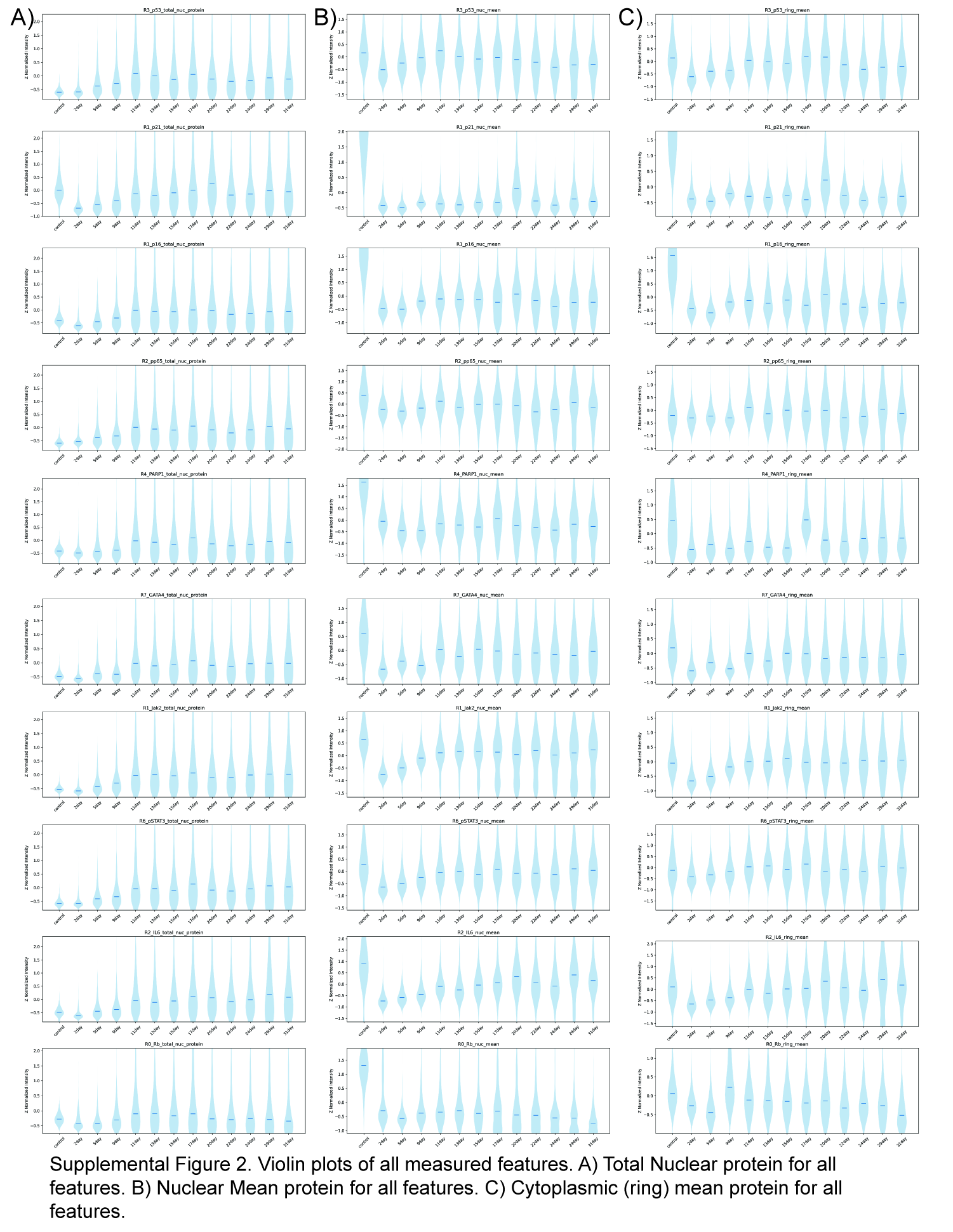

### Supplemental Figure 2.2

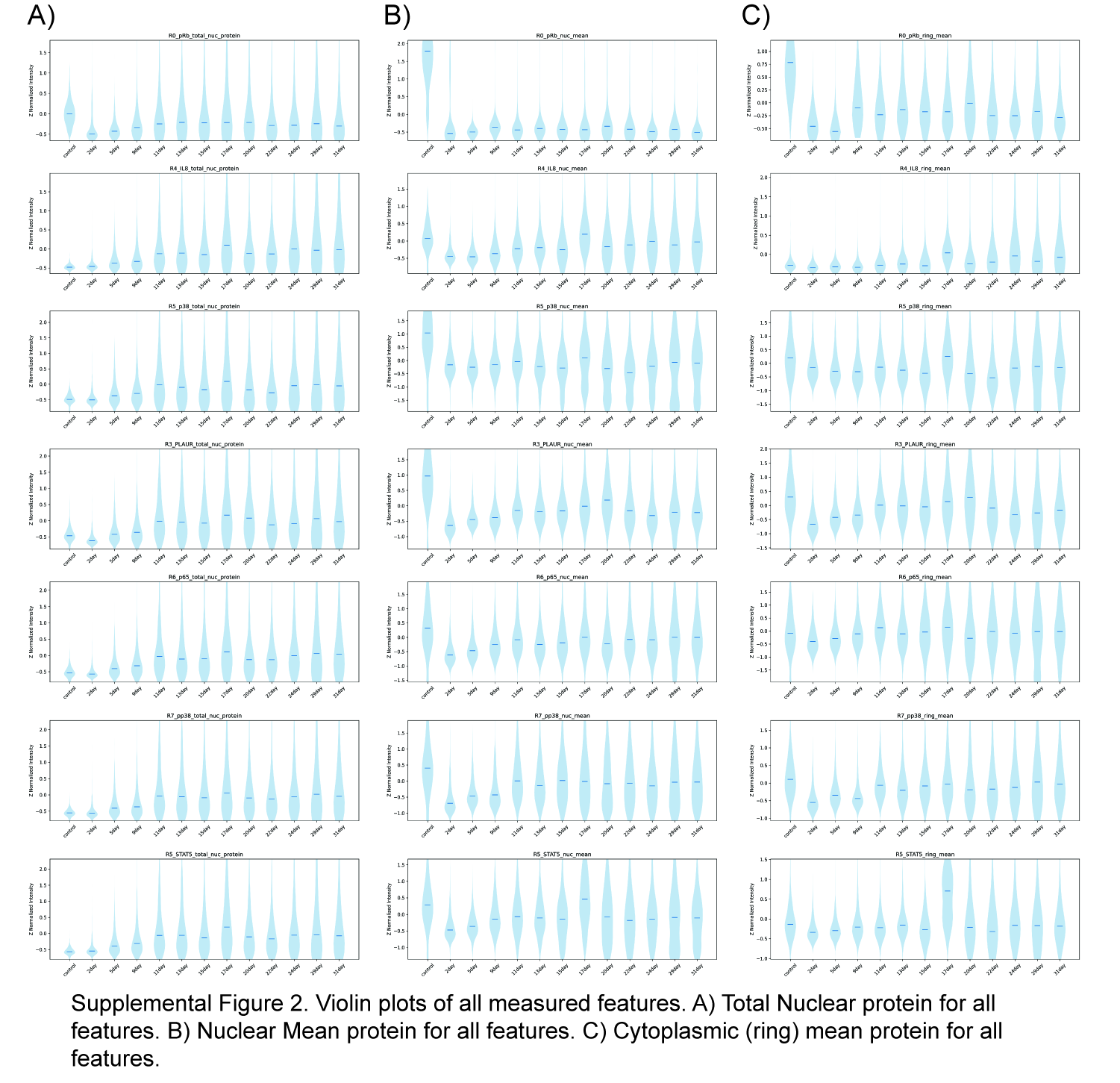

### Supplemental Figure 3

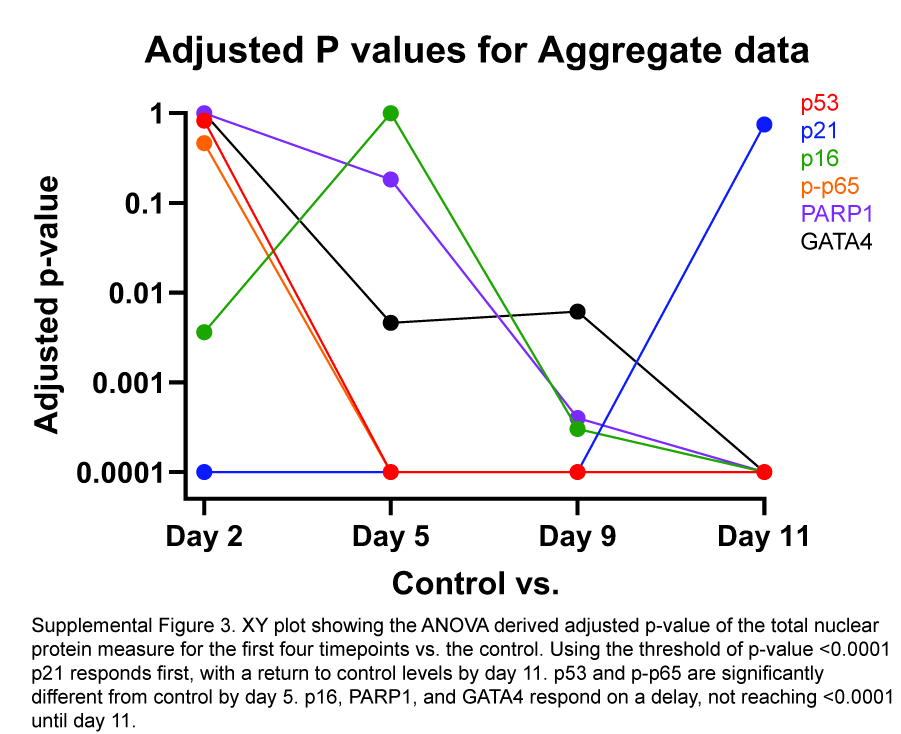

### Supplemental Table 1

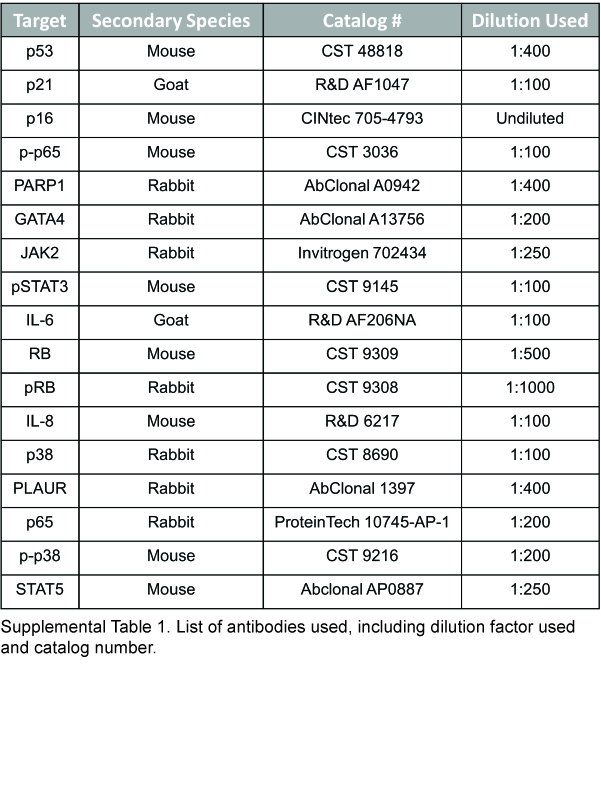
